# Low dose pig anti-influenza virus monoclonal antibodies reduce lung pathology but do not prevent virus shedding

**DOI:** 10.1101/2021.10.08.463636

**Authors:** Basudev Paudyal, Adam McNee, Pramila Rijal, B. Veronica Carr, Alejandro Nunez, John W. McCauley, Rodney S. Daniels, Alain R. Townsend, John A. Hammond, Elma Tchilian

## Abstract

We have established the pig, a large natural host animal for influenza, with many physiological similarities to humans, as a robust model for testing the therapeutic potential of monoclonal antibodies (mAbs). In this study we demonstrated that prophylactic intravenous administration of 15mg/kg of porcine mAb pb18, against the K160-163 site of the haemagglutinin, significantly reduced lung pathology and nasal virus shedding and eliminated virus from the lung of pigs following H1N1pdm09 challenge. When given at 1mg/kg, pb18 significantly reduced lung pathology and lung and BAL virus loads, but not nasal shedding. Similarly, when pb18 was given in combination with pb27, which recognised the K130 site, at 1mg/kg each, lung virus load and pathology were reduced, although without an apparent additive or synergistic effect. No evidence for mAb driven virus evolution was detected. These data indicate that intravenous administration of high doses was required to reduce nasal virus shedding, although this was inconsistent and seldom complete. In contrast the effect on lung pathology and lung virus load is consistent and is also seen at one log lower doses, strongly indicating that a lower dose might be sufficient to reduce severity of disease, but for prevention of transmission other measures would be needed.

## Introduction

Influenza virus infection remains a global health threat to humans and animal influenza A viruses are the CDC’s top zoonotic pathogen. Pig anatomy and physiology closely resembles that of humans. They have a similar distribution of sialic acid receptors in their respiratory tract and are infected with similar influenza A viruses, making them a powerful natural host large animal model to study immunity to influenza (1, 2). Monoclonal antibodies (mAbs) are promising therapeutics for virus infections including influenza (3-8). We have previously shown that the strongly neutralizing human IgG1 monoclonal antibody, 2-12C, administered prophylactically at 15mg/kg to pigs, reduced virus load and lung pathology after H1N1pdm09 challenge (9). However, the study could not be extended beyond 5-6 days due to the development of anti-human IgG responses. To overcome this limitation, we have isolated porcine mAbs from H1N1pdm09 infected pigs (10). The porcine antibodies were directed towards the two major immunodominant HA epitopes – the Sa site (residues 160 and 163) and Ca site (residue 130), also recognized by humans. The *in vitro* neutralizing activity of the pig mAbs was comparable to the strongest human mAbs. One of these mAbs, pb27, targeting the same HA1 site as 2-12C, encompassing residue K130, abolished lung and broncho-alveolar lavage (BAL) virus load and greatly reduced lung pathology after intravenous administration at 10mg/kg *in vivo*, although nasal shedding was not eliminated (10). Interestingly when administered at a lower dose of 1mg/kg, both human 2-12C and porcine pb27 reduced significantly lung pathology and lung virus load, suggesting that potentially the cost of therapy could be greatly reduced (9, 10).

Here we wished to determine the potency of another porcine mAb, pb18, against the HA1 site encompassing K160, recognised by many human sera and mAbs. We tested prophylactic administration at 15mg/kg and 1mg/kg. We also asked whether the effect of pb18 may be augmented additively or synergistically by the simultaneous administration of pb27, which targets the epitope encompassing K130.

## Materials and methods

### Ethics statement

Animal experiments were approved by the Pirbright institute ethics committee and Animal and Plant Health Agency (APHA) according to the UK animal (Scientific Procedures) Act 1986 under project licence P47CE0F2. Both Institutes conform to the ARRIVE guidelines.

### Monoclonal antibodies

Generation of porcine H1N1pdm09-specific mAbs were described previously (10). For animal studies, pb18 and pb27 were produced in bulk by Absolute Antibody Ltd (Redcar, UK) and dissolved in 25 mM Histidine, 150 mM NaCl, 0.02 % Tween, pH 6.0 diluent.

### Animal Studies

Twenty 5 weeks old, Landrace X Hampshire cross female pigs were obtained from a commercial high-health status herd and screened for antibody-free status against four swine influenza virus antigens: H1N1pdm09, H1N2, H3N2 and avian-like H1N1. Pigs weighed between 11-12kg (average 10kg). Pigs were randomized into four groups of five animals as follows: the first group received 15mg/kg of pb18; the second group received 1mg/kg of pb18; the third group received a combination of 1mg/kg of pb27 and 1mg/kg pb18 and; the fourth control group received PBS only. The mAbs were administered to the ear vein of animals sedated with stresnil (Janssen pharmaceuticals). Twenty-four hours after mAb(s) administration, all animals were challenged intranasally with 1×10^6^ PFUs of pandemic swine H1N1 isolate, A/swine/England/1353/2009 (H1N1pdm09) in 2ml (1ml per nostril) using a mucosal atomization device (MAD300; Wolfe Tory Medical). Clinical signs (temperature, loss of appetite, recumbence, skin haemorrhage, respiratory change, nasal discharge, altered behaviour) were observed and recorded. Clinical signs were mild and no animal developed moderate or severe disease.

### Gross Pathology and Histopathological scoring of Lung Lesions

At *post mortem*, the lungs were removed and digital photographs taken of the dorsal and ventral aspects. Lung tissue samples from the right cranial, middle, and caudal lung lobes were excised from the lung and collected into 10% neutral buffered formalin for routine histological processing. Formalin-fixed tissues were paraffin wax-embedded and 4µm sections cut and routinely stained with haematoxylin and eosin (H&E). Immunohistochemical detection of influenza A virus nucleoprotein (NP) was performed in 4µm tissue sections as previously described (11). Histopathological changes in the H&E-stained lung tissue sections were scored by a veterinary pathologist blinded to the treatment group. Lung histopathology was scored using five parameters (necrosis of the bronchiolar epithelium, airway inflammation, perivascular/bronchiolar cuffing, alveolar exudates and septal inflammation) scored on a 5-point scale of 0 to 4 and then summed to give a total slide score ranging from 0 to 20 per slide and a total animal score from 0 to 60 (12). The slides were also scored using the “Iowa” method, that also considers the amount of virus antigen present in the sample, as described (13).

### Tissue Sample processing

Two nasal swabs (one per nostril) were taken each day following challenge with H1N1pdm09. The swabs were placed into 2ml of virus transport medium (VTM) comprising tissue culture medium 199 (Sigma-Aldrich) supplemented with 25 mM 4-(2-hydroxyethyl)-1-piperazineethanesulfonic acid (HEPES), 0.035 % sodium bicarbonate, 0.5 % bovine serum albumin, penicillin 100 IU/ml, streptomycin 100 µg/ml, and nystatin 0.25 µg/ml, vortexed, centrifuged to remove debris, aliquoted and stored at ™80°C for subsequent virus titration. Serum samples were collected at the start of the study (prior to mAb administration and challenge) and at 0, 1, 3 and 4 DPI of challenge. Broncho-alveolar lavage fluid (BAL) was collected from the entire left lung with 150 ml of 0.1% BSA+PBS. BAL samples were centrifuged at 300 × *g* for 15 min, supernatant was removed, aliquoted, and frozen for antibody and virus titre analysis. The accessory lung lobes were dissected out and frozen at ™80°C for subsequent virus titration. The whole lobe was cut into pieces and 10% (w/v) pieces of lung were homogenized in 0.1% BSA using the gentle MACS Octo dissociator, the homogenate was clarified by centrifugation, and supernatant was used for virus titration.

### Virus titration

Virus titers in nasal swabs, BAL fluid and accessory lobe were determined by plaque assay on MDCK cells. The samples were 10-fold serially diluted in Dulbecco’s Modified Eagle’s Medium (DMEM) and 200µl overlaid on confluent MDCK cells in 12 well tissue culture plates. After 1 h, the wells were washed and overlaid with 1ml of 0.6% agarose containing culture medium with 2µg/ml of TPCK (Tosyl phenylalanyl chloromethyl ketone) trypsin. Plates were incubated for 48 h at 37°C. Overlay was removed, and plaques were visualized by staining the monolayer with 0.1% (v/v) crystal violet and enumerated.

### HA gene sequencing

Nasal swabs were collected in Trizol at 4 DPI and stored at -80°C. Subsequently, 0.4ml aliquots were thawed and extracted with 90µl of chloroform. Following centrifugation at 12,000 rpm/15min, the aqueous phase (∼250µl) was transferred to a 2ml Eppendorf tube and 1.5vol (375µl) 100% ethanol added, mixed by inversion, then centrifuged briefly. The liquid was transferred to a vRNA capture column (Qiagen: QIAamp viral RNA mini kit; #52906) and extracted following manufacturer’s instructions before eluting in 50µl of supplied AVE-buffer. RT-PCRs were performed using QIAGEN OneStep *ahead* RT-PCR kits (#220213) to amplify HA-gene products for Sanger sequencing and whole genome products for NGS (primer sequences available on request). Amplification products were then purified, sequenced, curated and analysed as described recently(14).

### Quantitation of mAbs in serum samples

Quantification of administered mAbs, pb18 and pb27, in serum, BAL and nasal swabs was determined by ELISA. Ninety-six well microtiter plates (Maxi Sorp, Nunc, sigma-Aldrich, UK) were coated with 1ug/ml of HA in PBS overnight at 4°C. Plates were blocked with 200µl of blocking solution composed of 4% milk powder in PBS, supplement with 0.05% Tween-20 (PBS-T) for 2 h at room temperature. A standard curve for pb18 mAb was prepared as 1:2 serial dilutions starting at 500ng/ml in dilution buffer and added in duplicate to the assay plate. Samples and standard were incubated for 1 h at room temperature. The plates were washed four times with PBS-T and incubated with detecting antibodies; polyclonal goat-anti pig IgG HRP (1:20,000) (Bio-Rad). Binding of Abs was detected by developing with 50µl/well 3,3’,5,5’-tetramethylbenzidine (TMB) high sensitivity substrate solution (Biolegend, UK) and stopping with 50µl 1M sulphuric acid. The plates were read at 450 and 630 nm with the Cytation3 Imaging Reader (Biotek). The mAbs concentrations in samples were interpolated from the standard curve using a sigmoidal four-parameter logistic curve fit for log of the concentration using GraphPad Prism 8.3.

### Microneutralization assay

Neutralising Antibody titres were determined in serum using a microneutralization (MN) assay. Briefly, viruses were diluted in virus growth medium (VGM; DMEM-Penicillin-streptomycin-0.1% BSA) and titrated to give plateau expression of NP in 3×10^4^ MDCK-SIAT1 cells after overnight infection in 96-well flat-bottomed plates. Serum was heat inactivated at 56°C for 30 min. Serum was diluted in 50µl of VGM starting at 1:4 and serially double diluted. 50µl of diluted virus was added to the serum and incubated for 1 h at 37°C. In each well 100µl of 3×10^4^ MDCK-SIAT1 cells were added and incubated overnight at 37°C. The monolayer was fixed with 4% paraformaldehyde and permeabilized with Triton-X100 and stained with mouse anti-NP IgG1 (clone AA5H, Bio-Rad Antibodies) followed by goat anti-mouse HRP (DAKO) antibody. TMB substrate was added and incubated for 2-5 min and reaction was stopped with 50µl of 1M sulphuric acid and absorbance was measured at 450 and 630nm (reference wavelength) on the Cytation3 Imaging Reader (Biotek). The MN titre was defined as the final dilution of serum that caused ≥50% reduction in NP expression.

### Statistical analysis

Non-parametric unpaired two-way t-test, two-way ANOVA multiple comparisons was performed using GraphPad Prism 8.3.

## Results

### mAb administration reduced virus load

To determine the *in vivo* efficacy of porcine mAbs, pb18 was administered at 15mg/kg, 1mg/kg or in combination with pb27 at 1mg/kg each (pb18+pb27). Controls were untreated animals. Twenty-four hours after mAb administration, all animals were challenged with H1N1pdm09 and culled at 4 days post infection (DPI) (**Fig. 1A**). The virus load in nasal swabs was assessed daily over the 4 days by plaque assay. Nasal virus loads were significantly lowered at 1, 2 and 3 DPI in the 15mg/kg pb18 group and when assessed by the area under the curve (AUC) compared to the control group (p=0.004) (**Figs. 1B and 1C)**. The AUC for the animals treated with combined pb18+pb27 at 1mg/kg was also reduced (p=0.056). However, 1mg/kg of pb18 alone was insufficient to reduce nasal virus shedding (p=0.126). We also assessed virus load in BAL and lung at 4 DPI (**Fig. 1D**). No virus was detected in the lungs of the 15mg/kg pb18 and pb18+pb27 treated groups at 4 DPI, while there was virus in 2 out of 5 pigs in the 1mg/kg pb18 group. Similarly, in BAL, virus load was significantly reduced in all mAb treated groups compared to controls. Overall, administration of pb18 at 15 mg/kg had a significant effect on virus shedding and lung and BAL virus loads, while the lower doses or combination of the two mAbs reduced lung virus and BAL loads. There was no evidence of synergistic effect of the pb18 plus pb27 cocktail.

**Figure 1:**
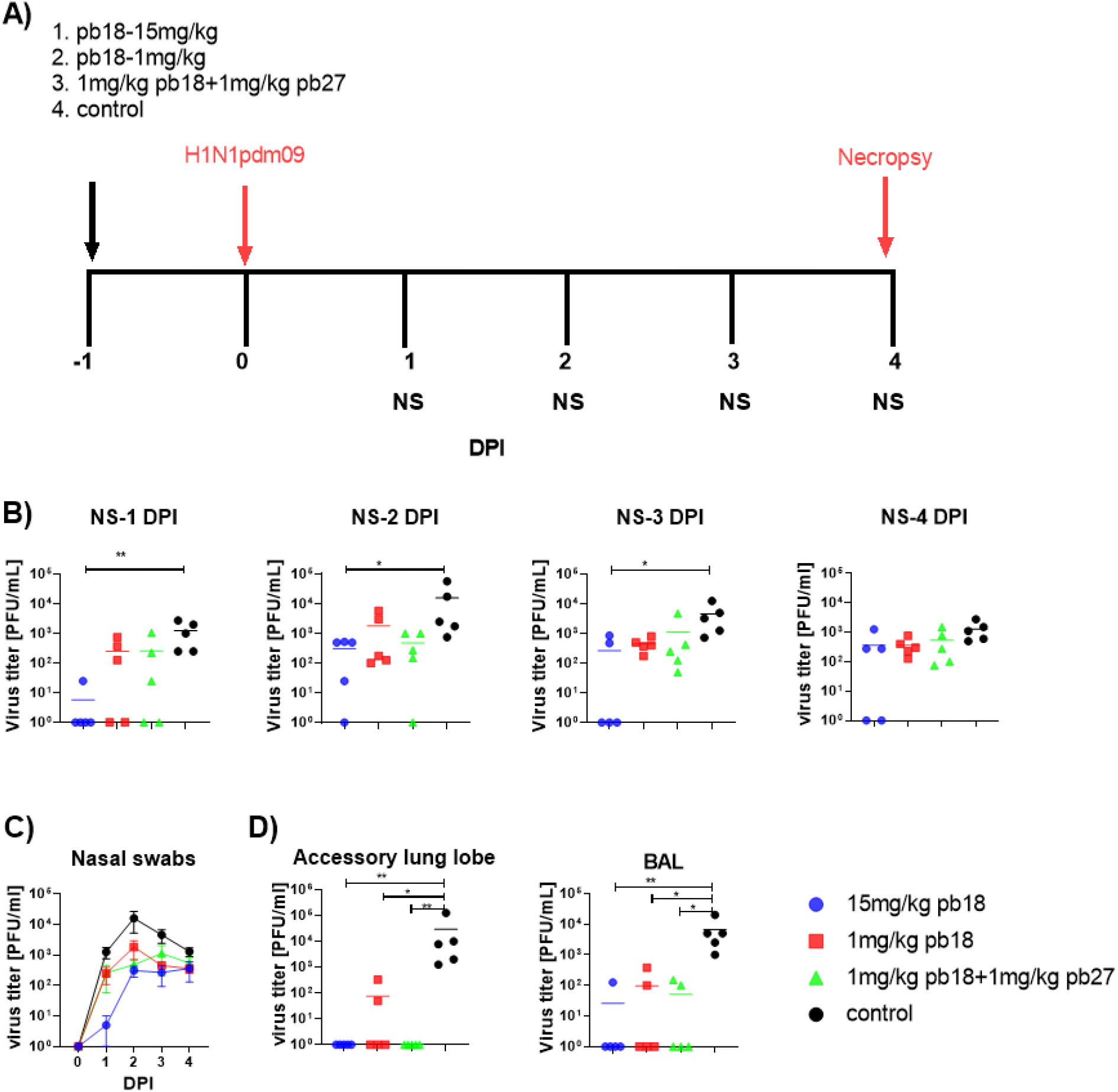
Experimental design and virus load. pb18 was administered at 15mg/kg, 1mg/kg or simultaneously with pb27 at 1mg/kg each to pigs which were challenged with H1N1pdm09 virus 24 hours later. The control group animals received PBS before infection. Nasal swabs were taken at 0, 1, 2, 3 and 4 DPI, and the pigs were culled at 4 DPI **(A)**. Virus load in daily NS **(B)** and over time **(C)**, accessory lung and BAL at 4 DPI (**D**) were determined by plaque assay. Each data point represents an individual within the indicated group and bars show SEM. The significance is indicated against the control. Asterisks denote significant differences *p<0.05, **p<0.01. **p<0.001, versus control as analysed by the one-way ANOVA Kruskal-Wallis test.

### mAb administration reduced lung pathology

Following challenge with H1N1pdm09, all control animals developed typical gross lung lesions indicative of influenza infection by day 4 DPI, as previously reported (15). In contrast the single 15mg/kg pb18 and combined pb18+pb27 at 1mg/kg treated animals showed significant (p=0.016) reductions in gross pathology (**Fig. 2A**). In control animals, characteristic histological lesions of influenza infection were observed, ranging from mild to severe necrotising bronchiolitis, with weakening of the bronchial and bronchiolar epithelium and neutrophilic exudation in bronchiolar lumina and alveoli. Areas of bronchointerstitial pneumonia with thickening of alveolar septa and peribronchial and perivascular infiltration by lymphohistiocytic cells were present. Immunolabelling for Influenza A nucleoprotein could be observed in bronchial and bronchiolar epithelium, macrophages in inflammatory exudates and occasional pneumocytes. The severity of these histopathological findings was assessed semi-quantitatively using the Iowa and Morgan methods (**Fig. 2B**). Compared to the control group, all mAb treated animals showed reduced severity of the pulmonary histopathological changes and lower numbers of influenza A nucleoprotein antigen immunolabelled cells (Fig 2B). Although reduction in histopathology (Morgan score) was observed in all mAb treated groups, this was significant (p=0.02) only in the 1mg/kg pb18 group. Significant reductions in Iowa score in the 15mg/kg pb18 (p=0.02) and 1mg/kg pb18 (0.008) groups but not in the combined pb18+pb27 mAb group were also detected. In contrast to the gross pathology results, there was an outlier for both Morgan and Iowa scores in the control group in which the histopathology scores were low. In this animal the gross lesions were mainly located in the left lung (sampled only for virological analysis) and no gross lesion was present in the histopathology specimen, resulting in a lower histology score and reduced statistical power when comparing scores with the treatment groups.

**Figure 2:**
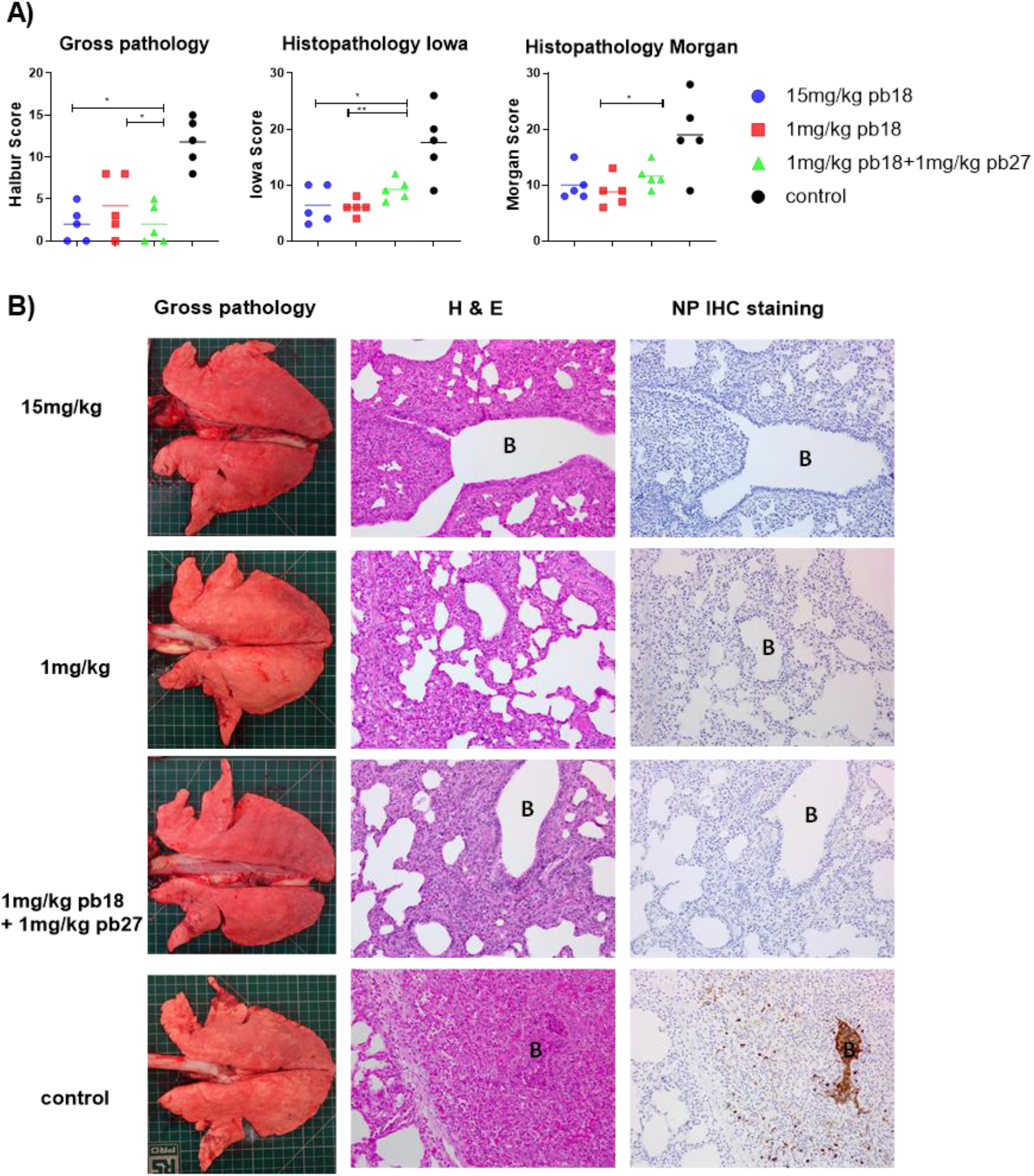
Lung pathology. pb18 was administered at 15mg/kg, 1mg/kg or simultaneously with pb27 at 1mg/kg each to pigs which were challenged with H1N1pdm09 virus 24 hours later. The control group animals received PBS before infection. The animals were culled four days later and the lung scored for appearance of gross and histopathology lesions. The gross and histopathology scores for each individual in a group and the group means are shown **(A)**. Representative gross pathology, histopathology (H&E staining; original magnification 100x), and immunohistochemical NP staining (original magnification 200x) for each group are shown (**B**), bronchiole region depicted at “B” the lesion scores and the histopathological scores (includes the NP staining). Pathology scores were analyzed using the one-way non-parametric ANOVA Kruskal-Wallis test. Asterisks denote significant differences *p<0.05, **p<0.01 versus control.

### Quantitation of mAbs in serum, BAL and Nasal Secretions (NS)

The concentration of administered mAbs in serum was determined by ELISA using recombinant HA glycoprotein at 0, 1, 3 and 4 DPI. Peak concentrations of 215μg/ml, 17μg/ml and 19μg/ml pb18 were detected in the 15mg/kg pb18, 1mg/kg pb18 and pb18+pb27 groups, respectively, at 24 h after administration. A decline in serum mAb concentrations was observed over the next 4 days to 101μg/ml, 9μg/ml, and 7μg/ml respectively (**Fig. 3A**). BAL samples showed the presence of administered mAbs; averages of 232ng/ml, 9ng/ml and 18ng/ml for 15 mg/kg pb18, 1 mg/kg pb18 and pb18+pb27 groups, respectively, were observed at 4 DPI. The administered mAbs were detected in nasal swabs at 4 DPI. We also analysed the neutralizing activity of the mAbs in serum and BAL. In serum, there were 50% inhibition neutralizing titers of 1:23,000, 1:1,200 and 1:2,900 for 15 mg/kg pb18, 1 mg/kg pb18 and pb18+pb27 groups, respectively, after 24 h of administration which gradually declined over the next four days (**Fig. 3B**). Microneutralization titers closely corresponded to the level of mAb in serum. Neutralizing activity in BAL was seen in all animals of the 15mg/kg pb18 group, in three animals administered 1mg/kg of both pb18 and pb27, but in no animals of the 1mg/kg pb18 group. The latter, probably relates to the lavage procedure which greatly dilutes the fluid present in airways.

**Figure 3:**
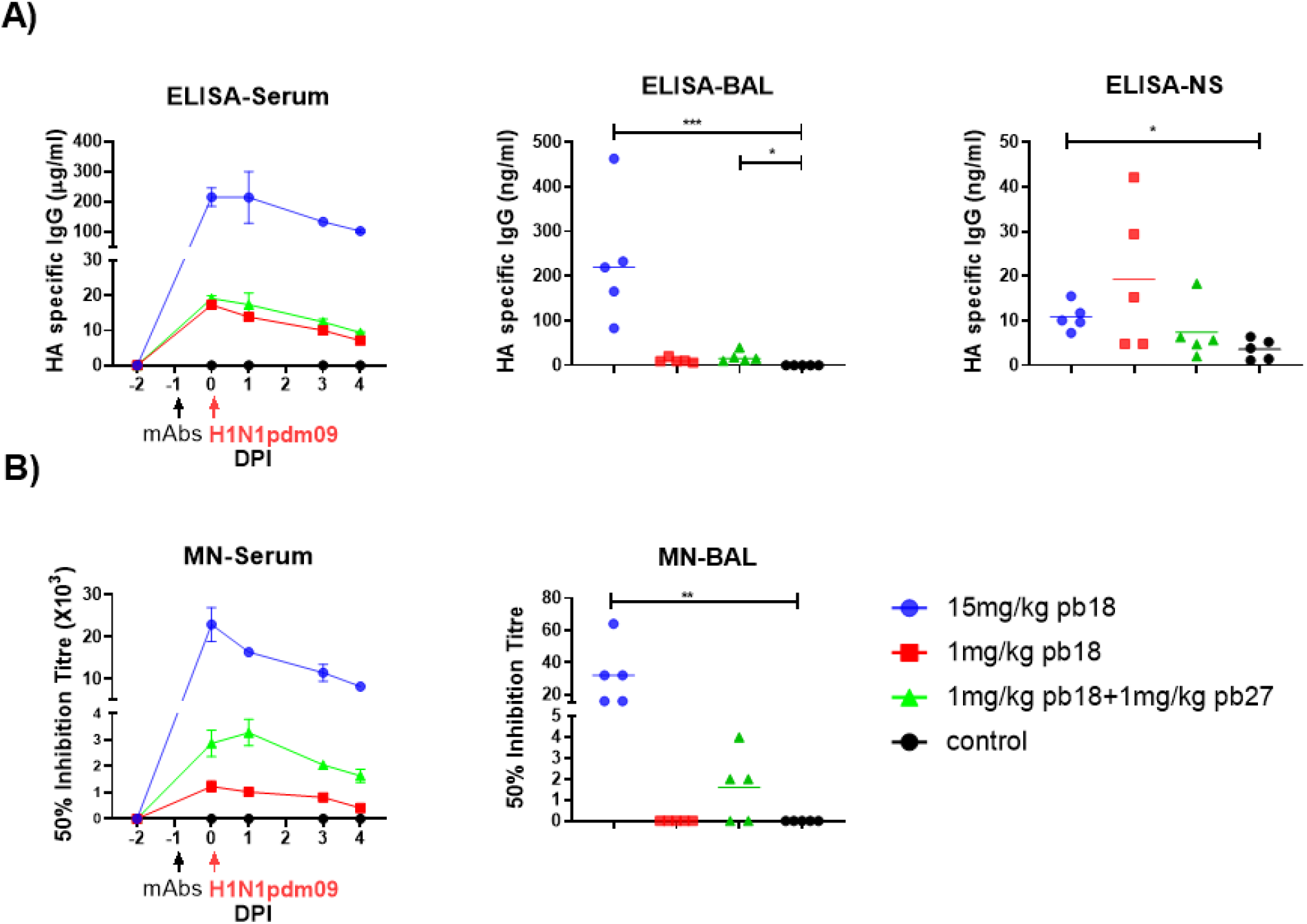
Concentration and neutralization titers of mAbs in serum, BAL and nasal swabs. HA specific IgG in serum was assessed by ELISA at the indicated DPI and in BAL and nasal swabs at 4 DPI (**A**). The 50% neutralization titers against H1N1pdm09 in sera at the indicated timepoints and in BAL at 4 DPI are shown (**B**). Symbols represent individual pigs within the indicated group and the lines represent SEM. Data were analyzed using the one-way non-parametric ANOVA Kruskal-Wallis test. Asterisks denote significant differences *p<0.05, **p<0.01, ***p<0.001 versus control.

**Figure 4:**
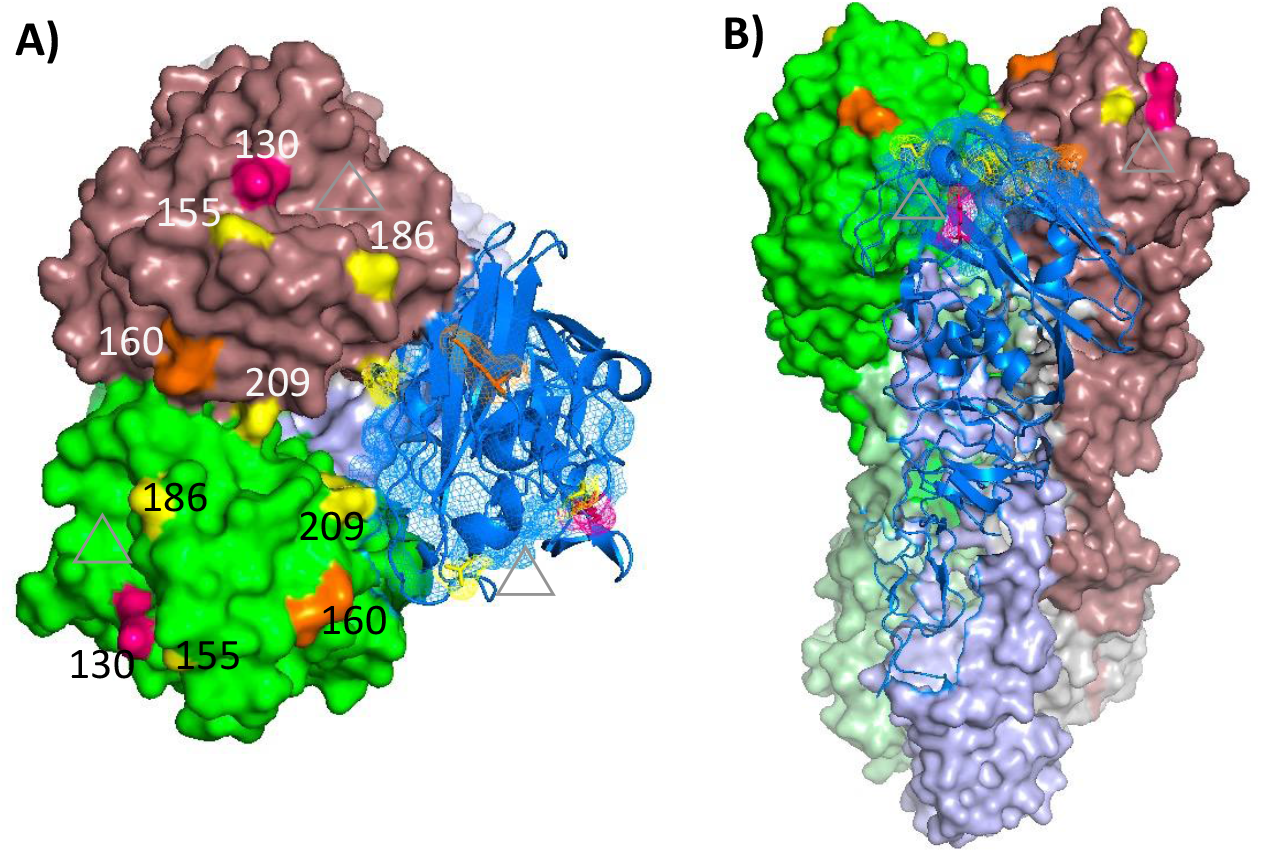
HA1 residues within the mAb binding sites. Residues that undergo substitution in mAb-escape variants are indicated: for pb27 K130 is indicated in pink and for pb18 K160 is indicated in orange. The HA amino acid variation in viruses (G155E, A186T and E209K) recovered from experimental animals are indicated in yellow. **A)** Top view, with sialic-acid binding sites indicated with gray triangles. **B)** Lateral view. In both views two monomers are shown in surface views, while the HA1 component of the third monomer gives a secondary structure view. Images were made using Pymol2 on PDB 4M4Y (45).

### Sequencing of virus

The stock of challenge virus and viruses from the nasal swabs of all 20 experimental animals were subjected to Sanger and whole genome sequencing with a focus on the HA gene. Complete HA gene sequences were recovered for the challenge virus and from 15 of the 20 animal specimens (**Table 1**). The challenge virus showed polymorphism at three positions in HA1, 154, 155 and 209. The dominant amino acid at these three positions (K154, G155 and E209) was non-polymorphic in in 15, 12 and 10 of the experimental animal specimens, respectively, and remained the dominant amino acid in all but one of these specimens. Animal 3194 in the 15mg/kg pb18 group showed dominance of E155 and K209, and A186T substitution. Two additional animals in the 15mg/kg pb18 group, 3190 and 3192, showed polymorphism at HA1 positions 184 and 186, or HA2 V201A substitution, respectively. One animal each in the 1mg/kg pb18, p18+p27 and the no mAb groups yielded sequences that showed polymorphism at HA1 position 222.

**Table 1:**
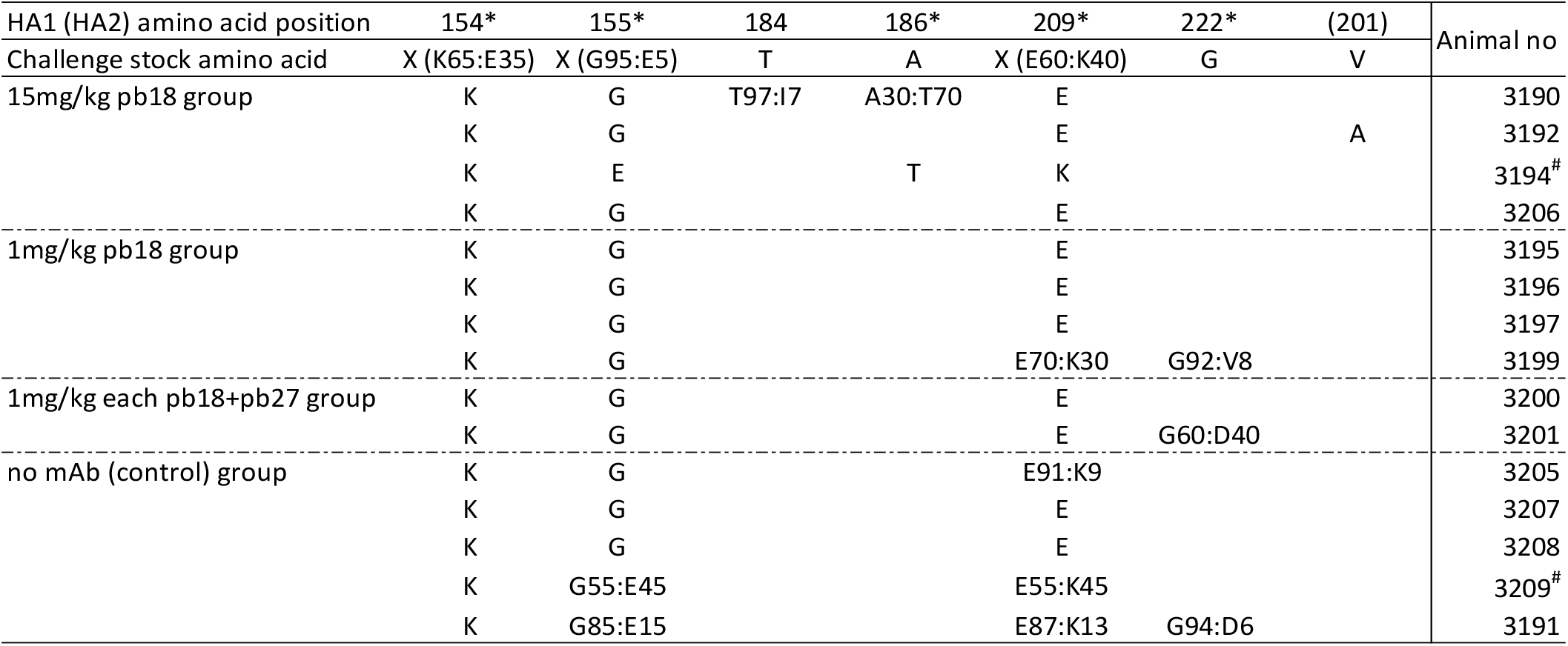
Virus HA amino acid variation in viruses recovered from experimental animals. Complete HA gene sequences were recovered for 15 of the experimental animals and the A/swine/England/1353/2009 challenge virus stock. Only positions that differ from the challenge virus are indicated. *Residues where amino acid polymorphism was seen in at least one virus. ^#^Sequences recovered by Sanger sequencing only where a cut-off of an amino acid representing ≥80% of the population was considered to represent no polymorphism (26).

| HA1 (HA2) amino acid position | 154* | 155* | 184 | 186* | 209* | 222* | (201) | Animal no |
| --- | --- | --- | --- | --- | --- | --- | --- | --- |
| Challenge stock amino acid | X (K65:E35) | X (G95:E5) | T | A | X (E60:K40) | G | V |  |
| 15mg/kg pb18 group | K | G | T97:I7 | A30:T70 | E |  |  | 3190 |
|  | K | G |  |  | E |  | A | 3192 |
|  | K | E |  | T | K |  |  | 3194<sup>#</sup> |
|  | K | G |  |  | E |  |  | 3206 |
| 1mg/kg pb18 group | K | G |  |  | E |  |  | 3195 |
|  | K | G |  |  | E |  |  | 3196 |
|  | K | G |  |  | E |  |  | 3197 |
|  | K | G |  |  | E70:K30 | G92:V8 |  | 3199 |
| 1mg/kg each pb18+pb27 group | K | G |  |  | E |  |  | 3200 |
|  | K | G |  |  | E | G60:D40 |  | 3201 |
| no mAb (control) group | K | G |  |  | E91:K9 |  |  | 3205 |
|  | K | G |  |  | E |  |  | 3207 |
|  | K | G |  |  | E |  |  | 3208 |
|  | K | G55:E45 |  |  | E55:K45 |  |  | 3209<sup>#</sup> |
|  | K | G85:E15 |  |  | E87:K13 | G94:D6 |  | 3191 |

## Discussion

In this study we demonstrated that prophylactic intravenous administration of porcine mAb pb18 at 15mg/kg significantly reduced lung pathology and nasal virus shedding, and abolished lung virus load in pigs following H1N1pdm09 challenge. When given at 1mg/kg, pb18 significantly reduced lung gross and histopathology and lung and BAL virus loads, but not nasal shedding of virus. Similarly, when pb18 was given in combination with pb27 at 1mg/kg each, lung virus load and pathology were reduced, although without an apparent additive or synergistic effect. The challenge virus showed amino acid polymorphism at three positions (154, 155 and 209) in HA1 and additional polymorphism or substitution was seen at positions 184, 186 or 222 in some of the samples from experimental animals. While there was no evidence for direct mAb driven evolution of the virus HA gene in any of the experimental animals, amino acid substitutions at the equivalents of all six of these positions have been associated with antigenic drift/mAb escape and/or changes in receptor specificity/host range in at least one influenza A HA subtype. Notably, substitutions at positions 155 (16-19), 186 (20-22), 209 (23) and 222 (24, 25) in the HA of A(H1N1)pdm09 viruses have been associated with such effects. The apparent lack of mAb driven HA evolution may partly be explained by the fact that samples were taken at only 4 DPI and tight bottlenecks may occur with very few viruses initiating infection (26). Such a scenario could explain the HA sequence derived from animal 3194 with HA1 G155E, A186T and E209K amino acid substitutions, the infecting virus being a minor variant within the A/swine/England/1353/2009 challenge virus stock.

The present data are in agreement with our previous experience with the human 2-12C and porcine pb27 mAbs in similar challenge experiments (9, 10). Intravenous administration of high doses (15mg/kg or 10 mg/kg) was required to reduce nasal virus shedding, although this was inconsistent and seldom complete, as in the present experiment. In contrast the effect on lung pathology and lung virus load is consistent and is also seen at one log lower doses in all experiments. This strongly indicates that a lower dose might be sufficient to reduce severity of disease, but for prevention of transmission other measures would be needed. It may be that a different IgG subclass or local administration to the respiratory tract would be more effective in supressing nasal virus shedding (27). Furthermore it is interesting to note that the effect of low dose mAb in reducing lung pathology, but not virus shedding is very similar to the effect of the powerful influenza specific CD8 responses induced by S-FLU immunisation (28, 29). This contrasts strongly with the effect of cytotoxic T cells in mice, which have been shown to protect both against disease (weight loss) and to reduce viral load (30-32), whereas in pigs a powerful T cell response or in this case mAb are insufficient to protect the upper respiratory tract from infection and shedding, although the lung viral load and pathology are reduced. This is an important difference between large and small animal models and it will be important be determine which best predicts the outcome of therapy in humans.

There was no evidence for a synergistic or additive effect for the pb18 and pb27 cocktail, although the mAbs targeted different epitopes. There is already a large literature on attempts to improve efficacy of mAbs by administering them as cocktails, which may result in lowered infection rates and virus loads, thereby reducing the probability of neutralization-escape variants emerging (33-36). While mAb cocktails alone have not proved effective in influenza, combination therapy with the viral polymerase inhibitor favipiravir and mAbs against the receptor-binding site and stem of virus HA completely stopped virus replication in nude mice, resulting in virus clearance (6, 37, 38). Administration of both anti-HA and anti-NA antibodies might also be effective and further studies to define whether the pigs generate broadly inhibiting anti-NA antibodies as has been shown in humans would allow us to test how protective these are *in vivo (39)*. High-resolution structures revealed a mechanism of cooperativity in Ebola virus (EBOV) mAb cocktails (40). A two-antibody cocktail offered protection in mice against the most antigenically divergent virus and demonstrated high therapeutic efficacy against live EBOV challenge in non-human primates. These findings offered a rational strategy for development of a potent two-antibody cocktail design based on structural features of mAb interactions with EBOV. However in other systems molecular interactions in which two or more mAbs recognize the same antigen synergistically are poorly defined, but these interactions might contribute greatly to the overall efficacy of protective mAb cocktails (41-44).

In summary we showed that the pig is a useful model to test mAb delivery and efficacy, particularly for influenzas virus, since pigs are infected by the same H1N1pdm09 influenza A viruses as humans and we have shown that pigs mount very similar antibody responses to the virus HA as seen in humans. The lobar and bronchial structure of the pig lung is very like that of humans and here we showed that a low dose of mAb may be highly effective in preventing lung pathology and severe disease.

## Acknowledgements

We are grateful to the veterinarians, pathologists and animal staff at APHA for providing excellent animal care and support during post-mortem sampling and to Helen Everett for her scientific support. We thank APHA for providing the swine A/Sw/Eng/1353/2009 influenza virus strain (DEFRA and devolved administrations of Scotland and Wales surveillance programme SV3041).

## Author contributions

ET, BP, AT, JH conceived, designed and coordinated the study. BP, AM, VC, performed animal experiments, processed samples and analyzed the data. AN carried out postmortem and pathological analyses; AT and PR designed experiments, provided reagents and developed assays for assessment of antibody function RD, JM performed sequencing analysis; ET, BP, PR and RD wrote and revised the manuscript.

